# CUT&Tag-BS: an efficient and low-cost method for simultaneous profiling of histone modification and DNA methylation

**DOI:** 10.1101/2021.07.19.452967

**Authors:** Ruifang Li, Sara A Grimm, Paul A Wade

## Abstract

It remains a challenge to decipher the complex relationship between DNA methylation, histone modification, and the underlying DNA sequence with limited input material. Here, we developed an efficient, low-input, and low-cost method for simultaneous profiling of genomic localization of histone modification and methylation status of the underlying DNA at single-base resolution from the same cells in a single experiment by integrating CUT&Tag with tagmentation-based bisulfite sequencing (CUT&Tag-BS). We demonstrated the validity of our method for both active and repressive histone modifications using 250,000 mouse ESCs. CUT&Tag-BS shows similar enrichment patterns of histone modification to those observed in non-bisulfite-treated control; it further reveals that H3K4me1-marked regions are mostly CpG-poor, lack methylation concordance, and exhibit prevalent DNA methylation heterogeneity among the cells. We anticipate that CUT&Tag-BS will be widely applied to directly address the genomic relationship between DNA methylation and histone modification, especially in low-input scenarios with precious biological samples.

## Introduction

DNA methylation and histone modification are two main mediators of epigenetic regulation, structurally and functionally coordinating with each other to regulate a multitude of biological processes on chromatin. A wide range of different modifications exist on the N-terminal tails of histones, including acetylation, methylation, phosphorylation and ubiquitylation (Bannister and Kouzarides, 2011), and their relationship with DNA methylation differs depending on the type and location of the modification (Fu et al., 2020; Xiao et al., 2012). The interdependent deposition and mutual exclusion of DNA methylation with different histone modifications generate complex chromatin modification patterns that demarcate distinct functional elements in the genome (Roadmap Epigenomics et al., 2015). Traditionally, the spatial relationship of DNA methylation with a given histone modification is determined by parallel genomic mapping of the two marks using whole genome bisulfite sequencing (WGBS) and ChIP-seq, followed by integrative overlap analysis of the profiles. However, by overlaying profiles from independent assays on different batches of cells, the traditional approach only permits correlative analysis and incurs prohibitive sequencing cost to obtain adequate depth of coverage for DNA methylation. To more directly and cost-effectively interrogate the interplay between DNA methylation and histone modification, BisChIP-seq (Statham et al., 2012) and ChIP-BS-seq (Brinkman et al., 2012) were developed to measure DNA methylation specifically from genomic regions marked with a given histone modification through bisulfite sequencing of ChIP-captured DNA (Kagey et al., 2010). Those two methods use a ligation-based bisulfite sequencing library preparation strategy, which requires a large amount of ChIP-ed DNA (∼ 100 ng); thus, they are not suitable for low-input samples. To reduce the amount of ChIP-ed DNA required, EpiMethylTag (Lhoumaud et al., 2019) was developed recently using tagmentaion-based library preparation strategy instead. Despite the differences in library construction, the three methods all rely on crosslinked ChIP to capture chromatin fragments associated with protein of interest. As a result, they suffer from the same limitations as ChIP, such as requiring large amounts of input material, suboptimal signal-to-noise ratio, and poor resolution. Moreover, formaldehyde crosslinking damages DNA (Do and Dobrovic, 2015), and incomplete reverse crosslinking interferes with bisulfite conversion thus confounding DNA methylation measurement (Wen et al., 2017).

CUT&Tag (Kaya-Okur et al., 2019) is a cutting-edge technique recently developed for genome-wide mapping of protein-DNA interactions. As an alternative to ChIP, CUT&Tag employs protein A-Tn5 (pA-Tn5) for antibody-guided in situ tagmentation of target-bound DNA in native cells. It has several advantages over ChIP, including faster and simpler workflow, lower input requirement, higher signal-to-noise ratio, and fewer sequencing reads needed. Tagmentation-based WGBS (T-WGBS) (Wang et al., 2013) utilizes Tn5 assembled with methylated adapter to tagment genomic DNA for subsequent bisulfite sequencing library preparation. It has a shorter workflow and requires less input DNA while exhibiting higher reproducibility and robustness than conventional ligation-based bisulfite sequencing. We reasoned that if conducting CUT&Tag experiment using pA-Tn5 loaded with methylated adapter adopted from T-WGBS, the resulting CUT&Tag-DNA can be further subjected to bisulfite sequencing following the same library preparation procedure as T-WGBS. CUT&Tag was thus adapted to couple with bisulfite sequencing (CUT&Tag-BS) for simultaneous profiling of histone modification and DNA methylation from the same cells. As a proof of concept, we performed H3K4me1- and H3K9me3-CUT&Tag-BS on mouse embryonic stem cells (mESCs) to demonstrate the utility of our method in directly addressing the relationships of DNA methylation with both active and repressive histone modifications at low cost.

## Result

### CUT&Tag-BS for simultaneous profiling of histone modification and DNA methylation

To enable simultaneous measurement of DNA methylation and histone modification on the same DNA molecule, we developed CUT&Tag-BS by integrating the CUT&Tag procedure with tagmentation-based bisulfite sequencing (Figure 1). In our method, CUT&Tag is employed to generate tagmented DNA in native cells specifically at genomic binding sites of a given histone modification using pA-Tn5 under the guidance of target-specific antibody, and the CUT&Tag-DNA is then subjected to tagmentation-based bisulfite sequencing, resulting in a base-resolution DNA methylation map at genomic regions bound by the given histone modification (Figure 4A). Our method utilizes pA-Tn5 loaded with methylated adapter (Table S1) to make the resultant CUT&Tag-DNA compatible with subsequent tagmentation-based bisulfite sequencing library preparation. The library preparation procedure includes an oligonucleotide replacement and gap repair step to covalently append methylated adapter sequences to each single strand of tagmented DNA fragments, followed by bisulfite conversion and PCR amplification. During the gap-repair step, unmethylated nucleotides are used to fill in the 9-base gap. Those bases (the first nine bases of read 2 and the last nine bases before the adapter on read 1) serve as an internal control to determine the efficiency of bisulfite conversion, but they must be excluded from downstream methylation analysis. As a combination of two techniques both characterized by low-input requirement and simple workflow, CUT&Tag-BS is thus suitable for efficient simultaneous profiling of histone modification and DNA methylation with low number of cells.

**Figure 1.**
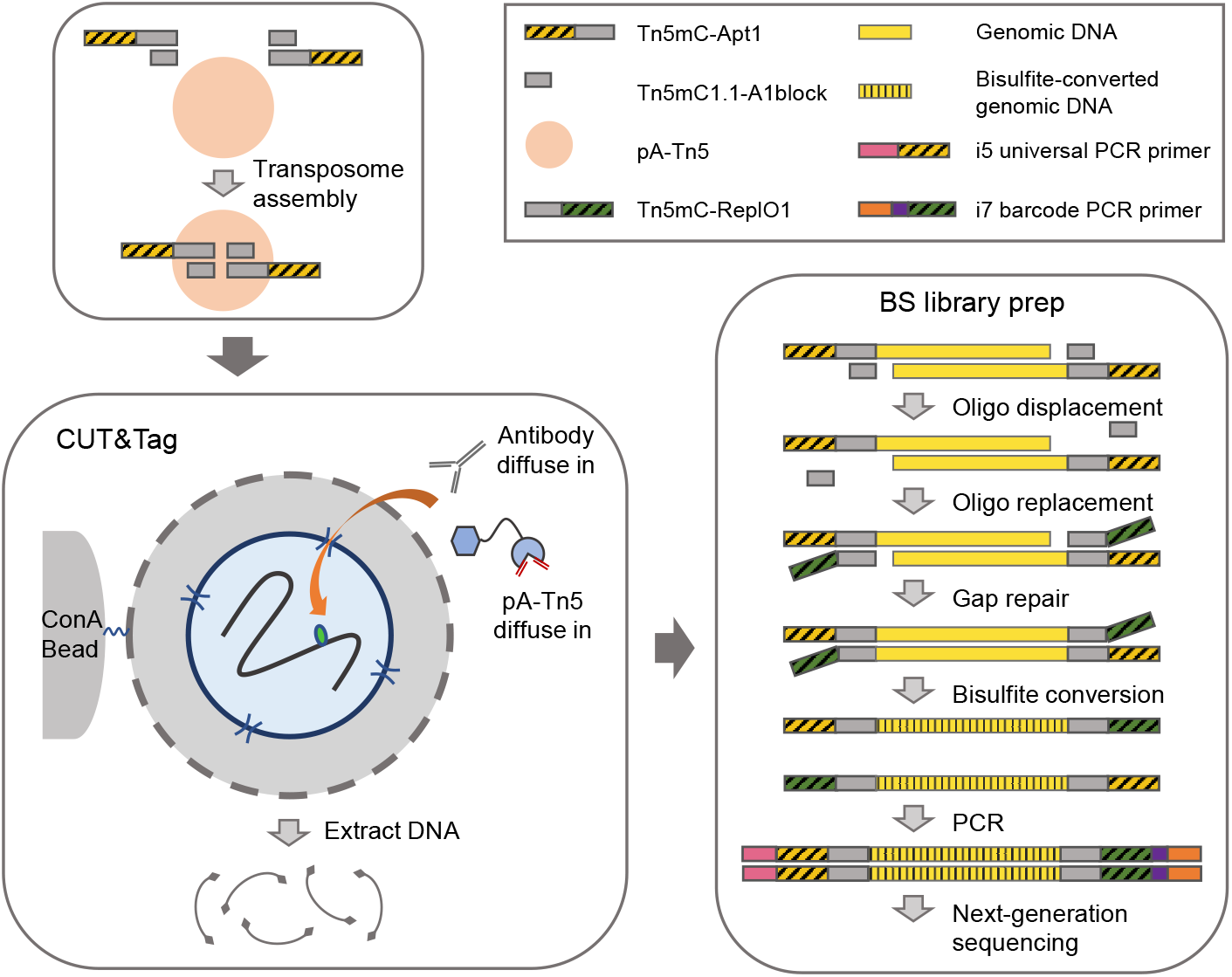
A schematic overview of CUT&Tag-BS workflow. CUT&Tag is adapted to couple with tagmentation-based bisulfite sequencing by using pA-Tn5 assembled with methylated adapter. The top adapter Tn5mC-Apt1 and the replacement oligonucleotide Tn5mC-ReplO1 must be methylated at all cytosines to maintain their identity during bisulfite treatment. CUT&Tag is performed according to published protocol (Kaya-Okur et al., 2019) with pA-Tn5 loaded with methylated adapter. Briefly, cells are harvested and bound to concanavalin A-coated magnetic beads. Cell membrane is permeabilized with digitonin (indicated by holes in the membrane) to allow the antibodies and pA-Tn5 to diffuse into the cells to find their targets and then tagment genomic binding sites of target protein. Tagmented DNA from CUT&Tag is subsequently subjected to tagmentation-based bisulfite sequencing library preparation. The oligonucleotide replacement and gap repair step covalently attaches the methylated adapter Tn5mC-ReplO1 to each DNA strand, followed by bisulfite conversion and PCR amplification to generate library for sequencing.

### CUT&Tag-BS shows comparable target enrichments to non-bisulfite-treated control

As a proof of principle, we performed CUT&Tag-BS for both active and repressive histone modifications (H3K4me1 and H3K9me3) using 250,000 mESCs. Since bisulfite treatment damages DNA extensively, to assess whether it alters CUT&Tag enrichment signals, we also generated non-bisulfite-treated CUT&Tag libraries as controls, which were sequenced and analyzed in parallel with CUT&Tag-BS libraries using paired-end 35-bp reads (Run-1). To determine the proper sequencing read length for this technique, CUT&Tag-BS libraries were further sequenced with paired-end 75-bp reads (Run-2). Similar to non-bisulfite-treated controls, CUT&Tag-BS libraries exhibited nucleosomal ladder pattern in fragment size distribution, although the proportion of smaller fragments increased after bisulfite treatment (Figure 2B), likely due to its detrimental effects on DNA integrity. Nevertheless, CUT&Tag-BS libraries showed similar alignment rates to corresponding non-bisulfite-treated controls, with ∼ 80% of H3K4me1 reads and ∼ 46% of H3K9me3 reads uniquely mapped to the reference genome (Figure 2A). Of note, H3K9me3 is enriched at repetitive regions (Martens et al., 2005), therefore, a relatively high percentage of H3K9me3 reads (∼ 37% of 35-bp reads; ∼ 24% of 75-bp reads) were ambiguously mapped (Figure 2A), leading to lower percentage of uniquely mapped reads. Lastly, CUT&Tag-BS libraries exhibited high bisulfite conversion rate (99.5%). Overall, based on library quality control metrics, all the CUT&Tag-BS libraries were of high quality irrespective of active or repressive histone modification being profiled.

**Figure 2.**
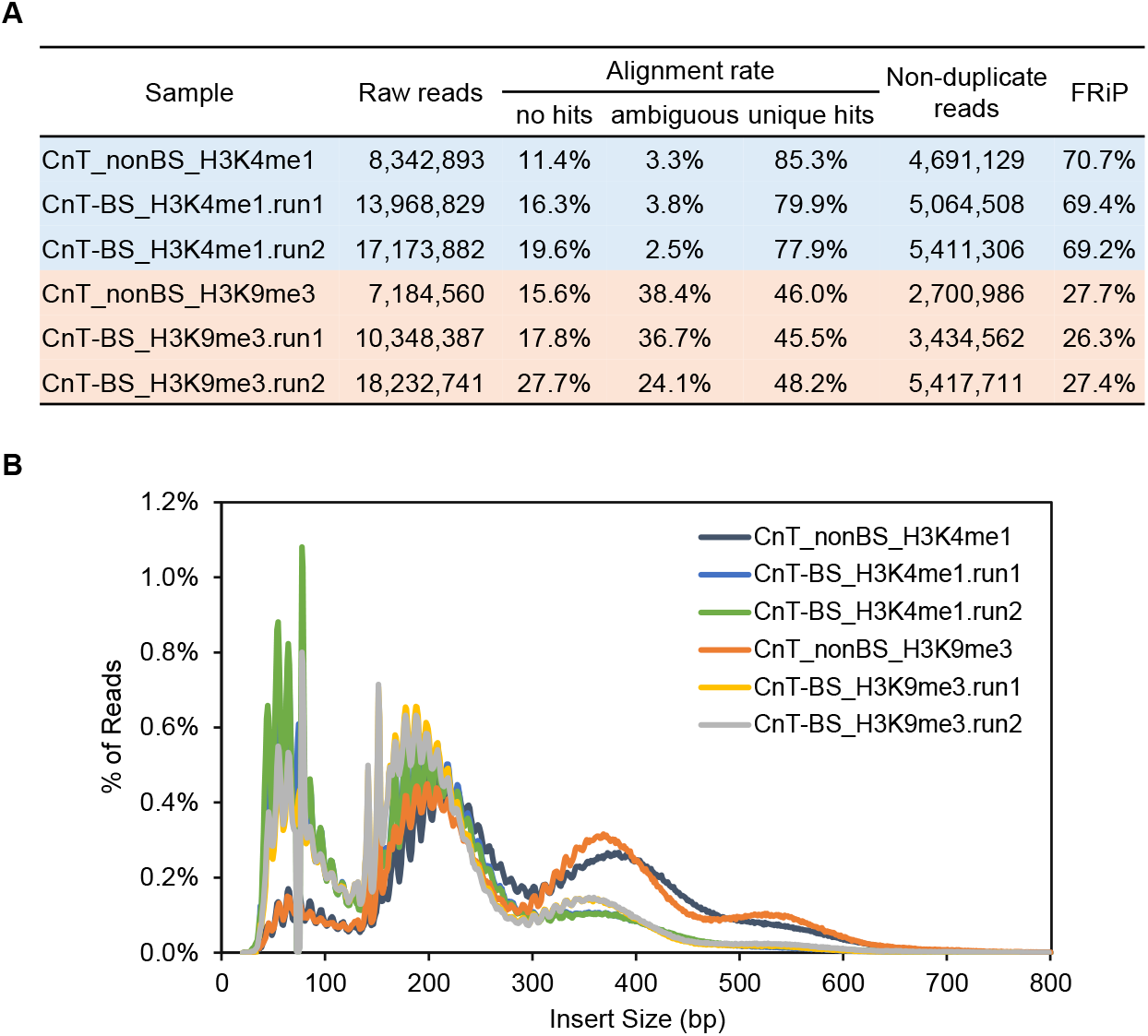
CUT&Tag-BS shows comparable library quality control metrics to non-bisulfite-treated control. (A and B) Sequencing metrics (A) and insert size distribution (B) of CUT&Tag-BS libraries compared with non-bisulfite-treated control libraries.

To assess whether bisulfite treatment distorts CUT&Tag enrichment signals, peak calling was performed on both CUT&Tag-BS and non-bisulfite-treated control samples. Despite bisulfite conversion, CUT&Tag-BS detected a similar number of peaks largely overlapping with those identified in the control sample (Figure 3A). Furthermore, the peak signals exhibited high correlation between CUT&Tag-BS and the corresponding control (Pearson’s *r* = 0.994 and 0.978 for H3K4me1 and H3K9me3, respectively) (Figure 3B). Moreover, CUT&Tag-BS showed similar FRiP score to that observed in corresponding control (Figure 2A), indicating comparable signal-to-noise ratio regardless of bisulfite conversion. Collectively, these analyses suggest that no obvious signal bias was observed after bisulfite treatment. Since different histone modifications are associated with distinct functional elements in the genome, to further verify that CUT&Tag-BS correctly identifies the expected functional elements, we conducted overlap analysis of detected peak regions with chromatin states of mESC defined by ChromHMM model (Pintacuda et al., 2017). H3K4me1-peaks were enriched preferentially at enhancers (∼ 55% peaks) and promoters (∼ 18% peaks) while H3K9me3-peaks were mainly located at heterochromatin (∼ 28% peaks) and intergenic regions (∼ 62% peaks) (Figure 3C), consistent with histone modification patterns used to define the chromatin states by ChIP-seq data. Taken together, CUT&Tag-BS achieved comparable enrichments of histone modification at expected genomic regions despite bisulfite treatment.

**Figure 3.**
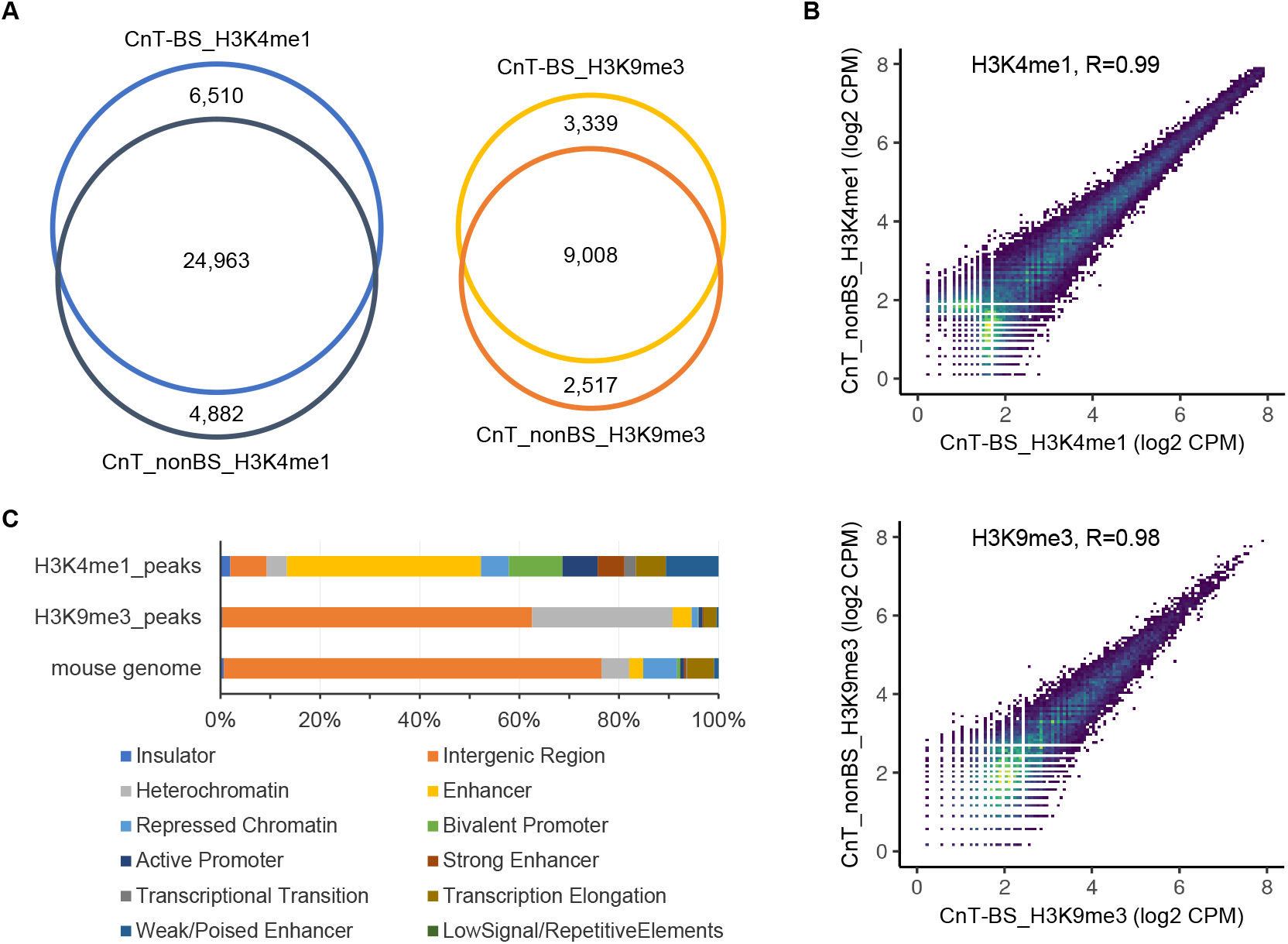
CUT&Tag-BS shows similar target enrichments to non-bisulfite-treated control. (A) Venn diagrams showing overlap of peaks identified in CUT&Tag-BS and corresponding non-bisulfite-treated control. (B) Density scatter plots displaying correlation of peak signals between CUT&Tag-BS and corresponding non-bisulfite-treated control. Each dot represents an individual peak in the unified peak set with viridis color scale indicating density. Pearson’s *r* value is shown at the top of each plot. (C) Bar graph showing the proportion of detected peaks and the reference mouse genome falling into each chromatin state of mESC.

**Figure 4.**
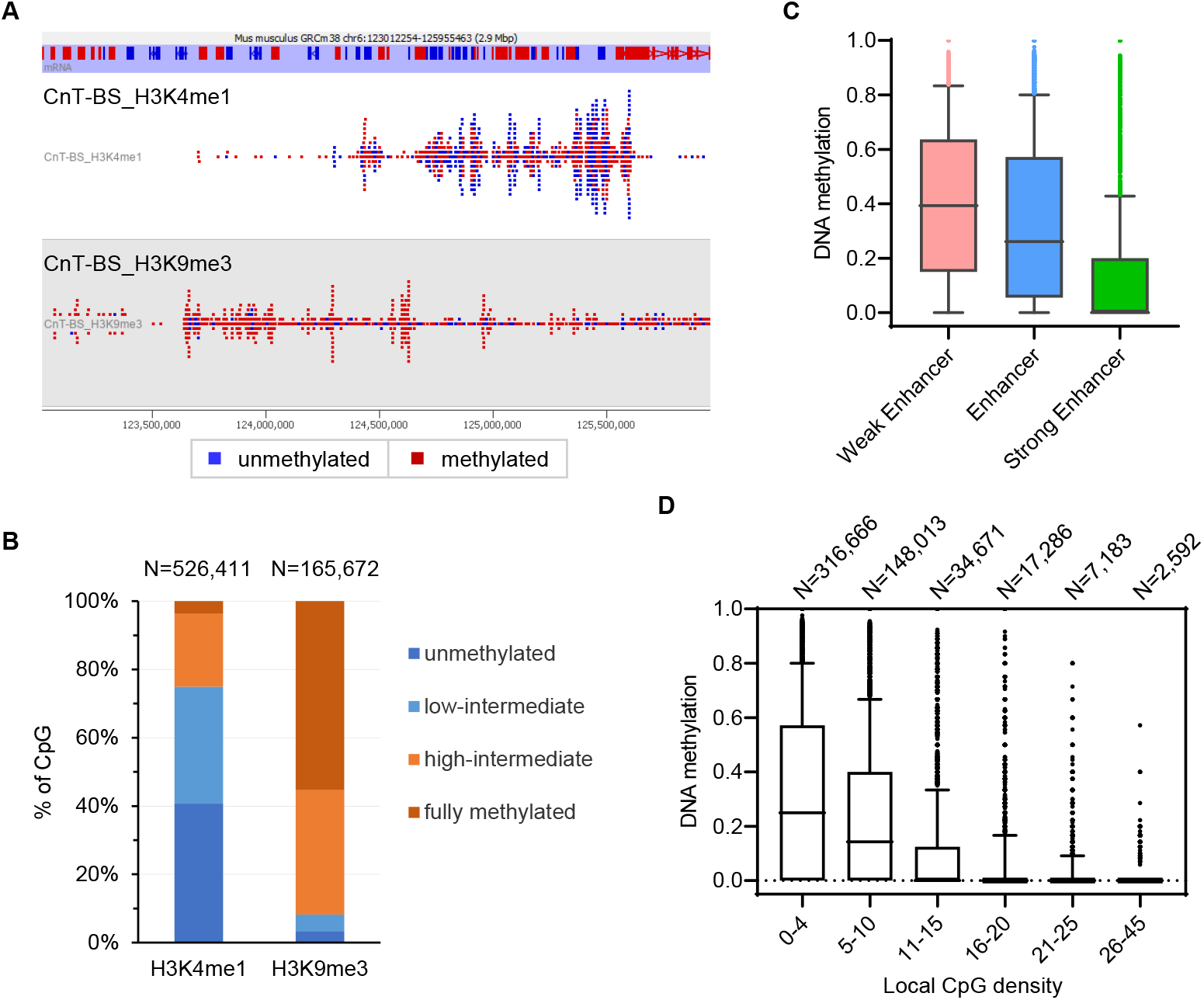
CUT&Tag-BS simultaneously measures DNA methylation at target binding sites. Example of CUT&Tag-BS tracks at chr6:123,012,254-125,955,463 with CpG methylation status indicated by color (red: methylated; blue: unmethylated). (B) Bar graph showing the percentage of H3K4me1- or H3K9me3-marked CpGs in each methylation category (unmethylated: < 10% methylation; low-intermediate: 10-50% methylation; high-intermediate: 50-90% methylation; fully methylated: > 90% methylation). (C) Box-and-whisker plot showing methylation levels of H3K4me1-marked CpGs (minimum of 5× coverage) at different types of enhancers. The whiskers represent the 10^th^ and 90^th^ percentiles. (D) Box-and-whisker plot showing methylation levels of H3K4me1-marked CpGs (minimum of 5× coverage) with different local CpG density, which is defined as the number of CpG sites within region ± 100 bp of a given CpG. The whiskers represent the 10^th^ and 90^th^ percentiles. The number of CpGs in each group is shown at the top of the plot.

### CUT&Tag-BS simultaneously measures DNA methylation at target binding sites

In addition to histone modification enrichments, CUT&Tag-BS simultaneously measures DNA methylation at H3K4me1- and H3K9me3-marked genomic regions (Figure 4A). M-bias plots show an average CpG methylation rate of ∼ 32.5% for H3K4me1-reads and ∼ 81.5% for H3K9me3-reads across the read length, except for the first nine bases of read 2 (Figure S1A), which correspond to the 9-bp gap region filled in with unmethylated nucleotides. Since a CpG site is either methylated or unmethylated in a given single cell, when DNA methylation is profiled from a cell population, the methylation level at a given CpG site actually reflects the percentage of cells methylated at that site with intermediate methylation indicating methylation heterogeneity among the cells. The composite methylation level averages across all profiled CpG sites; thus, the observed intermediate methylation level may result from averaging across CpGs with different methylation levels or may simply represent a similar intermediate methylation state across all profiled CpG sites. To ascertain which is the case, we further investigated the distribution of methylation levels of individual CpG sites in peak regions. Irrespective of the minimum coverage required at the CpG sites, we observed consistent skewed distributions of CpG methylation levels with H3K4me1-marked CpGs peaking at 0% methylation while H3K9me3-marked CpGs peaking at 100% methylation; however, a substantial number of CpGs exhibited intermediate methylation levels in both cases (Figure S1B). To quantitatively determine the proportion of CpGs with different methylation status, we classified CpG methylation levels into four categories, less than 10% methylation (unmethylated), 10-50% methylation (low-intermediate), 50-90% methylation (high-intermediate), and more than 90% methylation (fully methylated). CpG sites covered by at least five reads at peak regions were then categorized based on their methylation levels. Among H3K9me3-marked CpGs, 55.3% were fully methylated, 36.4% exhibited high-intermediate methylation, and the rest showed low-intermediate methylation (4.8%) or were unmethylated (3.4%) (Figure 4B), consistent with previous notion that H3K9me3 is associated with hypermethylated DNA in mESCs (Brinkman et al., 2012). In contrast, 40.7% of H3K4me1-marked CpGs were unmethylated, 34.3% showed low-intermediate methylation, 21.5% exhibited high-intermediate methylation, and 3.5% were fully methylated (Figure 4B). It suggests that even though H3K4me1-marked histones generally bind to unmethylated DNA, they also bind to methylated DNA at some genomic locations in a subset of the cells as reflected by intermediately methylated H3K4me1-CpGs, in line with previously reported cell-to-cell DNA methylation heterogeneity at enhancers in mESCs (Angermueller et al., 2016; Song et al., 2019).

H3K4me1 is a chromatin hallmark of enhancers (Bulger and Groudine, 2011), however, the role of DNA methylation at enhancers is poorly understood. We thus investigated DNA methylation at different types of enhancers. Although ‘Weak/Poised Enhancer’, ‘Enhancer’, and ‘Strong Enhancer’ were all marked with H3K4me1, they showed different DNA methylation levels diminishing gradually with the increase of enhancer activity (Figure 4C), suggesting negative association of DNA methylation with enhancer activity in general. However, DNA methylation was highly variable at ‘Weak/Poised Enhancer’ and ‘Enhancer’ (Figure 4C). To determine other influencing factors of DNA methylation at H3K4me1-marked CpG sites, we further investigated the relationship between DNA methylation status and local CpG density around the site. H3K4me1-marked CpGs were mostly located at low-CpG-density regions, and their methylation levels were inversely correlated with the local CpG density (Figure 4D), in concordance with previously observed global conflicts of CpG density and DNA methylation in human and mouse tissues (Chen et al., 2018).

### CUT&Tag-BS reveals lack of methylation concordance at enhancers

To determine the proper read length for CUT&Tag-BS, we first explored the necessity of sequencing through the library insert. We reasoned that if CpG methylation status is consistent across the insert, it would be valid to use CpGs present on the partially sequenced portion to represent all CpGs on a given insert; otherwise, reading through the insert would be necessary to obtain more accurate information on DNA methylation at target-binding sites. We thus assessed concordance of methylation calls across the sequencing read containing three or more CpGs. More than one-third of the assessed H3K4me1- and H3K9me3-reads (42.1% and 33.7%, respectively) showed mixed methylation (Figure 5A), indicating inconsistent methylation status of CpGs on the same insert DNA. It is seemingly contrary to the previous findings that DNA methylation levels were strongly correlated at nearby CpG sites particularly when they are within 1 to 2 Kb from each other (Bell et al., 2011; Eckhardt et al., 2006). However, our analysis of CpG methylation concordance within single sequencing reads only assessed one allele of a given cell, while previous findings were based on CpG methylation levels of cell population. To rule out the possibility that the discrepancy was due to methodology difference, we further measured the correlation and difference of methylation levels between pairs of neighboring CpG sites as a function of their genomic distance. Correlation of methylation levels rapidly dropped below 0.5 within _2_00 bp of distance for both H3K4me1- and H3K9me3-marked CpGs (Figure 5B). However, the absolute methylation difference between neighboring CpGs exhibited different patterns in the two marks; H3K9me3-marked CpGs showed a constant small difference in methylation levels at neighboring CpGs irrespective of their distance, while H3K4me1-marked CpGs displayed larger methylation differences that positively correlated with genomic distance between the sites (Figure 5B), suggesting that methylation concordance is highly dependent on the genomic regions interrogated. Nevertheless, considering that longer reads greatly increased CpG coverage (Figure 6B), sequencing through the library insert would be beneficial for achieving higher accuracy in DNA methylation. Therefore, the proper read length for CUT&Tag-BS is subject to the library insert size pertinent to the target protein.

**Figure 5.**
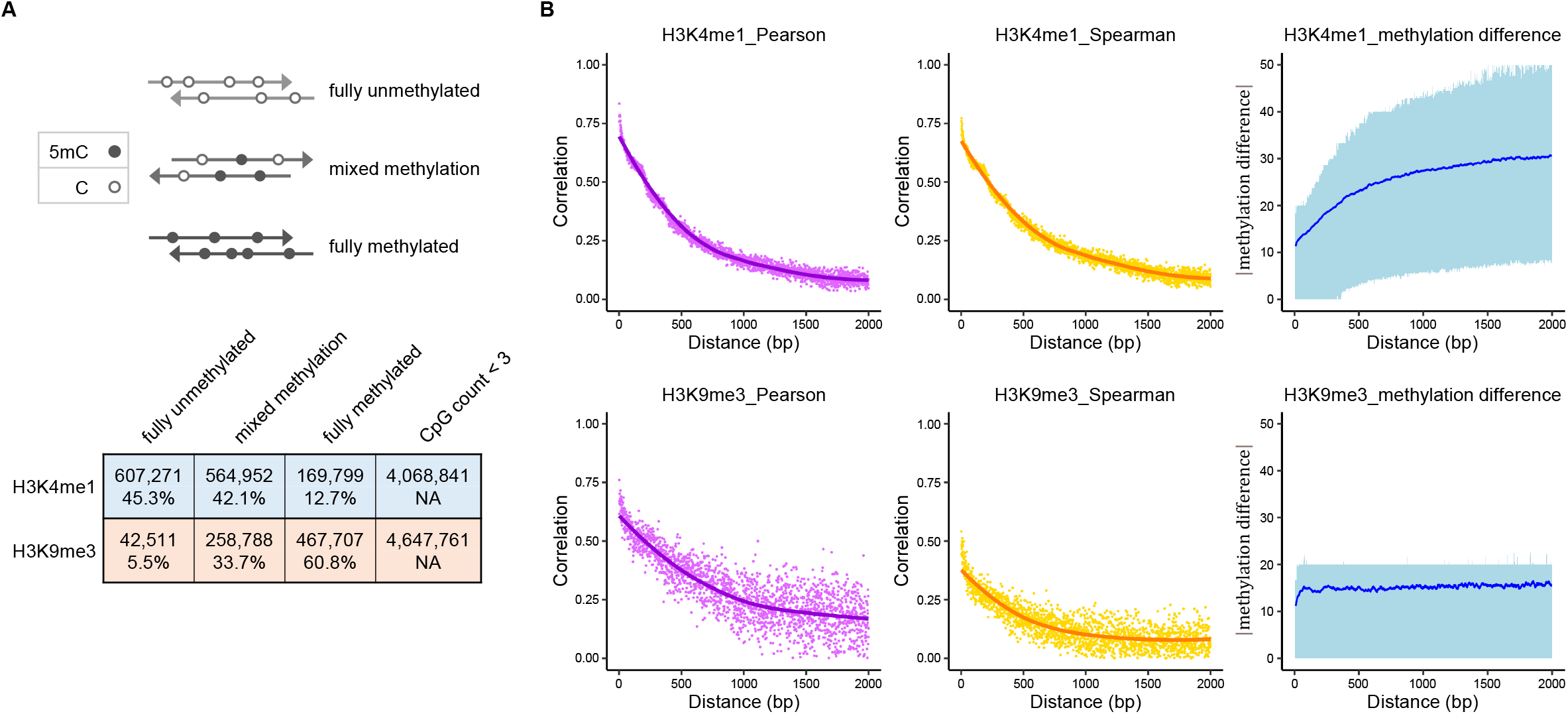
CUT&Tag-BS reveals lack of methylation concordance at enhancers. (A) Concordance of CpG methylation calls on individual sequencing reads of CUT&Tag-BS. Only reads with three or more CpGs were assessed. The ‘lower’ and ‘upper’ thresholds were set at 10% and 90% methylation, respectively, to split the reads into three categories (fully unmethylated, mixed methylation, and fully methylated). (B) Correlation (Pearson: left panel; Spearman: middle panel) and difference (right panel) of methylation level between neighboring CpG sites as a function of their genomic distance. The x-axis represents genomic distance between pairs of CpG sites, with at least one CpG per pair located in the peak regions (H3K4me1: top panel; H3K9me3: bottom panel). Loess curves were added to the correlation plots (left and middle panels) in ggplot2 v3.3.2 with geom_smooth. The absolute methylation differences between neighboring CpGs are shown (right panel), with the dark blue line and the light blue ribbon indicating the average and interquartile range, respectively.

**Figure 6.**
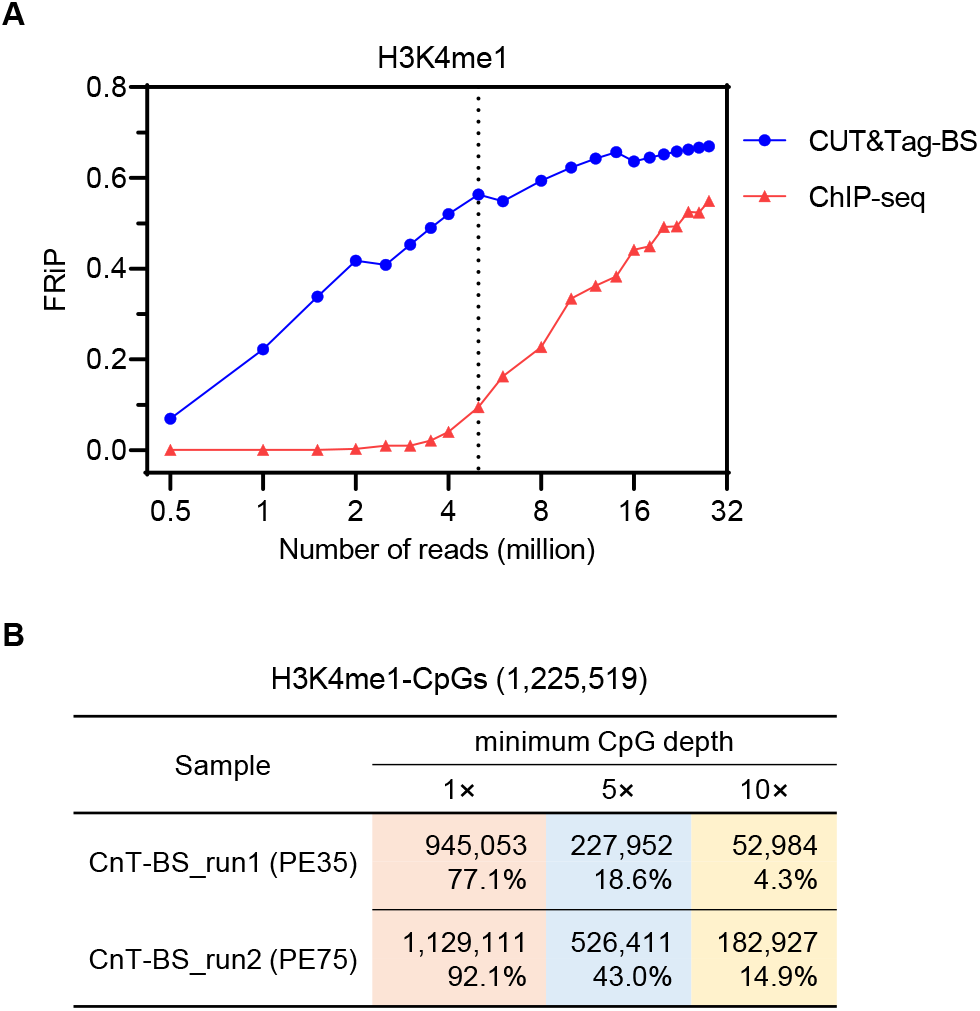
CUT&Tag-BS is a low-cost method requiring fewer sequencing reads. (A) FRiP scores calculated based on HOMER peak calls using different numbers of randomly downsampled sequencing reads in CUT&Tag-BS compared with ChIP-seq. The same H3K4me1 antibody was used in the two assays on mESCs. The dashed line marks five million reads. (B) CpG coverage at H3K4me1-peaks when using different sequencing read lengths (PE35 vs. PE75).

### CUT&Tag-BS is a low-cost method requiring fewer sequencing reads

Lastly, we assessed the sequencing depth required for CUT&Tag-BS. Owing to its high signal-to-noise ratio, CUT&Tag needs only several million sequencing reads (Kaya-Okur et al., 2019). To verify that the advantage also extends to CUT&Tag-BS, we calculated FRiP score as a measure of signal-to-noise ratio, comparing H3K4me1-CUT&Tag-BS data with publicly available ChIP-seq data of H3K4me1 in mESCs (Feldmann et al., 2020). The FRiP score of CUT&Tag-BS nearly reached a plateau (∼ 0.6) with just 5 million sequencing reads (Figure 6A), similar to the number of reads claimed to be sufficient in CUT&Tag for histone modifications. In contrast, ChIP-seq data was less suitable for peak-calling when down-sampled to 5 million reads as indicated by the much lower FRiP score (∼ 0.1) (Figure 6A). Due to its intrinsic high background, ChIP-seq required ∼ 30 million reads to reach similar FRiP score as CUT&Tag-BS with only 5 million reads (Figure 6A). Therefore, CUT&Tag-BS allows one to reach adequate coverage of CpGs at target-binding sites with a greatly reduced amount of sequencing. For example, with 5.4 million usable reads (paired-end 75-bp) in H3K4me1-CUT&Tag-BS, 92.1% and 43.0% of the CpGs at H3K4me1-peaks were covered with a minimum depth of 1× and 5×, respectively (Figure 6B). In conclusion, CUT&Tag-BS shows higher signal-to-noise ratio than ChIP-seq with the same antibody, and thus is inherently more cost-effective than methods relying on ChIP to capture chromatin fragments associated with the histone modification of interest.

## Discussion

We developed CUT&Tag-BS to simultaneously profile genomic localization of histone modification and methylation status of the underlying DNA from the same cells by coupling CUT&Tag with bisulfite sequencing (Figure 1). The conventional bisulfite sequencing strategy requires relatively large amounts of high-quality DNA for optimal conversion (Clark et al., 1994; Clark et al., 2006), thus imposing challenges on the methods that rely on crosslinked ChIP to capture chromatin fragments for subsequent bisulfite sequencing. CUT&Tag-BS overcomes some of the limitations of existing methods. First, CUT&Tag-BS uses native cells without formaldehyde fixation thus avoiding the side effects of crosslinking/decrosslinking, as reflected by high bisulfite conversion efficiency (99.5%) and improved overall alignment rate in CUT&Tag-BS (∼80%) (Figure 2A) compared with EpiMethylTag (∼65%) and ChIP-BS-seq (∼33%) (Lhoumaud et al., 2019). Second, CUT&Tag-BS adopts a tagmentation-based bisulfite sequencing library preparation strategy, which is faster and requires substantially less DNA than conventional ligation-based bisulfite sequencing library construction (Wang et al., 2013). This, in combination with efficient CUT&Tag, makes CUT&Tag-BS suitable for much lower cell numbers than is practical with existing methods relying on ChIP. Third, CUT&Tag-BS exhibits high signal-to-noise ratio (Figure 6A). The inherent low background and high read mappability translate to almost an order-of-magnitude reduction in the amount of sequencing required, thus dramatically lowering sequencing cost. In addition, low-background CUT&Tag-BS ensures DNA methylation measurements derive from DNA truly bound by the given histone modification, thus providing more confidence in the observed relationship of the marks than using methods relying on high-background ChIP.

Notably, bisulfite treatment did not alter the pattern of CUT&Tag enrichment in terms of FRiP score, number of detected peaks, genomic locations of the peaks, and signal strength at peaks (Figure 2A and 3). In addition to well-preserved enrichment signals, CUT&Tag-BS simultaneously measured DNA methylation at enriched regions with single-base resolution. Prevalent DNA methylation heterogeneity was observed at enhancers among the cells, as shown by the intermediate methylation levels of more than half (55.8%) of the H3K4me1-CpG sites in mESCs (Figure 4B). Since cell-to-cell differences are always present to some degree in any population of cells, potential epigenetic heterogeneity should be considered when deciphering the relationship between DNA methylation and histone modification in a cell population. Direct bisulfite sequencing of CUT&Tag-DNA enables more accurate and sensitive analysis of the spatial relationship between the marks than the traditional correlative approach, which neglects cell heterogeneity due to integrating data from independent assays on different cells. DNA methylation was previously reported to be characterized by spatial correlation within 1 to 2 Kb (Bell et al., 2011; Eckhardt et al., 2006), providing a rationale for assigning the methylation state of single-CpG measurements to all CpGs within a genomic interval. However, CUT&Tag-BS analysis indicated that DNA methylation concordance was highly dependent on the genomic regions interrogated, with H3K4me1-marked regions lacking methylation concordance (Figure 5), in line with recently reported faster methylation concordance decay at enhancers (Hui et al., 2018). In contrast to previously oversimplified relationship of the marks as either coexisting or mutually exclusive, CUT&Tag-BS revealed more complex relationship between DNA methylation and histone modification, which remains to be further elucidated in a region-specific manner.

In summary, CUT&Tag-BS was successfully developed with H3K4me1 and H3K9me3 as the histone modification of interest. It is straightforward and requires minimal adaptation of previous CUT&Tag (Kaya-Okur et al., 2019) and T-WGBS (Wang et al., 2013) protocols. As an efficient and low-cost method, CUT&Tag-BS needs only a small number of input cells and sequencing reads. Hence, we anticipate that CUT&Tag-BS will be widely applied to directly interrogate the complex interplay between DNA methylation and histone modification, especially in low-input scenarios with precious biological samples. Moreover, we speculate that CUT&Tag-BS could be used to investigate the relationship between DNA methylation and other important epigenetic regulators, such as chromatin modifiers/remodelers and transcription factors. By simultaneously providing information of both chromatin context and strand-specific DNA methylation at base resolution, CUT&Tag-BS would enable better understanding of epigenetic regulation in various biological contexts.

## Methods

### Assemble pA-Tn5 with methylated adapter

Single-stranded methylated adapter oligos (Tn5mC-Apt1 and Tn5mC1.1-A1block; Table S1) were resuspended in Annealing Buffer (10 mM Tris-HCl pH 8.0, 50 mM NaCl, 1 mM EDTA) to a final concentration of 200 µM. The resuspended oligos were mixed in equal volumes then annealed in a thermocycler using the following program; 95°C for 2 min, followed by decreasing the temperature in 5°C increments to reach 25°C with a ramp rate of -0.1°C/sec and 5 min incubation at each ending, then hold at 8°C. For transposome assembly, 100 μL of 8 µM pA-Tn5 fusion protein was mixed with 20 μL of the annealed adapter, and the mixture was incubated at room temperature (RT) for 1 hour on a rotator then stored at -20°C.

### CUT&Tag-BS

#### CUT&Tag

H3K4me1- and H3K9me3-CUT&Tag were performed on 250,000 mESCs using pA-Tn5 loaded with methylated adapter, following the CUT&Tag protocol (Kaya-Okur et al., 2019) with minor modifications. Harvested mESCs were counted with an automated cell counter. One half million of the cells were transferred into a new 1.5mL Eppendorf tube then pelleted at 600 × g for 3 min at RT. The cell pellet was washed twice with 1 ml Wash Buffer (20 mM HEPES pH 7.5, 150 mM NaCl, 0.5 mM Spermidine (Sigma-Aldrich), 1× Protease inhibitor cocktail (Sigma-Aldrich)), then resuspended in 200 μL of Wash Buffer and split equally into two tubes. For the cells to bind to Concanavalin A-coated magnetic beads (Bangs Laboratories), 10 µL activated beads were added into each sample and the cell-bead mixture was rotated at RT for 10 min. The tubes were then placed on a magnet stand to clear and the supernatant was removed. The bead-bound cells were resuspended in 100 µL of Antibody Buffer (20 mM HEPES pH 7.5, 150 mM NaCl, 0.5 mM Spermidine, 1× Protease inhibitor cocktail, 0.01% Digitonin (EMD Millipore), 2 mM EDTA, 0.1% BSA (NEB)) with anti-H3K4me1 antibody (Cell Signaling Technology) or anti-H3K9me3 antibody (Abcam) at 1:100 dilution. Primary antibody incubation was performed at 4°C overnight on a rotator. The tubes were then placed on the magnet stand to clear and unbound primary antibody was removed by discarding the supernatant. Next, the bead-bound cells were resuspended in 100 µL of Antibody Buffer with Guinea Pig anti-Rabbit IgG antibody (Antibodies online) at 1:100 dilution and then incubated at RT for 30 min on a rotator. After secondary antibody binding, the cells on beads were washed three times with 1 mL Dig-Wash Buffer (20 mM HEPES pH 7.5, 150 mM NaCl, 0.5 mM Spermidine, 1× Protease inhibitor cocktail, 0.01% Digitonin) to remove unbound antibodies. The bead-bound cells were then incubated with pA-Tn5 loaded with methylated adapter at 1:200 dilution in 100 µL Dig-300 Buffer (20 mM HEPES pH 7.5, 300 mM NaCl, 0.5 mM Spermidine, 1× Protease inhibitor cocktail, 0.01% Digitonin) for 1 hour at RT on a rotator. After pA-Tn5 binding, the cells on beads were washed three times with 1 mL Dig-300 Buffer to remove unbound pA-Tn5. To activate pA-Tn5 for tagmentation, the bead-bound cells were resuspended in 100 µL Tagmentation Buffer (20 mM HEPES pH 7.5, 300 mM NaCl, 0.5 mM Spermidine, 1× Protease inhibitor cocktail, 0.01% Digitonin, 10 mM MgCl_2_) then incubated at 37°C for 1 hour in a thermomixer with shaking at 1000 rpm. The tagmentation reaction was terminated by mixing with 500 µL Buffer PB from MinElute PCR Purification Kit (Qiagen). The tubes were then placed on the magnet stand to clear, and the supernatant was transferred into the MinElute column (Qiagen) to purify CUT&Tag-DNA according to the manufacturer’s instructions.

#### Oligonucleotide replacement and gap repair

At this step, the shorter adapter oligonucleotide, which is not covalently linked to the genomic DNA, is replaced by a methylated oligonucleotide (Tn5mC-ReplO1; Table S1), and the nine-base gap is repaired by combined actions of DNA polymerase and DNA ligase. Briefly, 11 μL purified CUT&Tag-DNA was used in the reaction (11 μL DNA, 2 μL 10 μM Tn5mC-ReplO1 oligo, 2 μL 10× Ampligase buffer (Lucigen), 2 μL dNTP mix (2.5mM each) (Thermo Fisher Scientific)) assembled in a PCR tube. The reaction was first incubated in a PCR thermocycler with the following program; 50°C for 1 min, 45°C for 10 min, ramp down to 37°C at a rate of - 0.1°C/sec, then hold at 37°C. Once the program reached 37°C, while the tubes remained in the thermocycler, 1 μL T4 DNA polymerase (NEB) and 2.5 μL Ampligase (Lucigen) were added into each sample separately. The reaction was mixed by pipetting up and down with a P20 micropipette then incubated at 37°C for 30 min. The reaction was stopped by adding 1 μL 0.5 M EDTA (pH = 8.0) then cleaned up with MinElute PCR Purification Kit (Qiagen) following the manufacturer’s instructions. The gap-repaired DNA was eluted in 22 μL Buffer EB, and 2 μL of the eluted DNA was saved as a control to assess DNA damage by bisulfite treatment.

#### Bisulfite conversion

The purified gap-repaired DNA was bisulfite converted with EZ DNA Methylation-Lightning Kit (Zymo Research) according to the manufacturer’s instructions. Briefly, 20 μL gap-repaired DNA and 130 μL Lightning Conversion Reagent were mixed then split equally into two PCR tubes (75 μL/tube). The reaction was incubated in a thermocycler as follows: 98°C for 8 min, 54°C for 60 min, then hold at 4°C. Subsequent purification and desulfonation was performed exactly as described in the user manual. Bisulfite converted DNA was eluted in 25 μL M-elution Buffer.

#### PCR amplification

Bisulfite converted DNA was amplified and barcoded in 50 µL PCR reaction (22 µL bisulfite-converted DNA, 25 µL 2× KAPA HiFi HotStart Uracil+ ReadyMix (Roche), 1.5 µL 10 µM i5 universal PCR primer, 1.5 µL 10 µM i7 barcode PCR primer) with the following PCR program: 98°C for 45 sec; 14 cycles of 98°C for 15 sec, 63°C for 30 sec, 72°C for 30 sec; final extension at 72°C for 2 min; then hold at 4°C. The reserved 2 μL control DNA (without bisulfite conversion) was amplified and barcoded in parallel using the same PCR program in 50 μL PCR reaction (2 μL gap-repaired DNA, 25 μL 2× NEBNext High-Fidelity PCR Master Mix (NEB), 1.5 μL 10 μM i5 universal PCR primer, 1.5 μL 10 μM i7 barcode PCR primer, 20 μL Nuclease-free H2O). The PCR reactions were cleaned up with AMPure XP beads (Beckman Coulter).

#### Sequencing

The libraries were quantified with Qubit dsDNA HS Assay Kit (Invitrogen) then pooled and sequenced on Illumina NextSeq550 with paired-end 35-bp or 75-bp reads.

### CUT&Tag-BS data analysis

Raw sequencing reads were filtered to remove any read pairs with mean base quality score less than 20 at either read end. Adapters were removed via Cutadapt v1.12 (Martin, 2011) with parameters “-a CTGTCTCTTATACAC -A CTGTCTCTTATACAC -O 5 -q 0 -m 20 -p”. For any read pairs in which adapter was identified and removed, an additional nine bases were trimmed from the 3’ end of read1, as they correspond to the 9-base gap region filled in with unmethylated nucleotides. Genomic alignment was performed by Bismark v0.23.0 (Krueger and Andrews, 2011) with parameters “-X 1000 --non_bs_mm” (all other parameters as default) against the mm10 reference assembly (GRCm38) with Bowtie2 v2.3.0 (Langmead and Salzberg, 2012) as the underlying mapper. Positional methylation bias information was reported by the Bismark v0.23.0 bismark_methylation_extractor tool with parameters “-p --include_overlap -- mbias_only”. Duplicate mapped fragments were removed by the Bismark v0.23.0 deduplicate_bismark tool with parameter “-p”. The observed insert size distribution per library was determined by Picard tools v1.110 CollectInsertSizeMetrics.jar (http://broadinstitute.github.io/picard). Per-residue methylation data was collected by the Bismark v0.23.0 bismark_methylation_extractor tool with parameters “-p --ignore_r2 9 -- comprehensive --mbias_off --bedGraph --cytosine_report”. The bisulfite conversion rate per library was calculated by assessing methylated and unmethylated cytosine counts at the first 9 bases of read2.

### Peak calling and peak overlap between samples

HOMER v4.10.3 (Heinz et al., 2010) was used to call peaks per sample as follows. Mapped read data was prepared using the makeTagDirectory function with parameters “-format sam -read1 - fragLength 200”, followed by peak calls using the findPeaks function with parameters “-size 500 -minDist 1000 -L 0 -region” for H3K4me1 and parameters “-size 1000 -minDist 2500 -L 0 - region” for H3K9me3. Peaks overlapping mm10 blacklist regions (https://github.com/Boyle-Lab/Blacklist/tree/master/lists) were discarded. For each histone mark, a unified peak set was generated by BEDtools v2.29.2 merge (Quinlan and Hall, 2010) as the union of peak calls from the CUT&Tag-BS library and the corresponding non-bisulfite-treated control. Analysis of shared peaks between samples was performed with subsets of unified peaks that overlapped the HOMER peak calls from given samples. Briefly, the unified peak set was intersected individually with HOMER peak calls from each sample in comparison, and the subset of unified peaks in the intersection was used to determine peak overlap between samples.

### Peak signal quantification

To quantify signal intensity at peaks, the mapped paired-end hits were converted to a single fragment via BEDtools v2.29.2 bamtobed with the “-bedpe” option. The number of fragments overlapping each unified peak was collected via BEDtools v2.29.2 coverage with the “-counts” option, and then subsequently converted to CPM (counts per million).

### Overlap of peaks with chromatin states

Chromatin states of mESC (Pintacuda et al., 2017) as defined by ChromHMM were downloaded from https://github.com/guifengwei/ChromHMM_mESC_mm10. Overlap of the detected peaks with chromatin states was determined by BEDtools v2.29.2 intersect. In cases of peaks overlapping multiple chromatin states, each peak was assigned to the chromatin state with which it showed the most overlap.

### Methylation concordance and methylation correlation

Methylation concordance across reads was calculated by summing up methylated and unmethylated cytosine counts per fragment identifier from the CpG context output file generated by the Bismark v0.23.0 bismark_methylation_extractor tool.

To assess correlation of CpG methylation as a function of genomic distance, all pairs of CpG sites within 2Kb were identified for which at least one CpG was within the unified peak regions of the given histone mark. The set of CpG pairs was further filtered to retain only those with at least 5× coverage for both sites. Pearson and Spearman correlations of methylation level between pairs of CpGs at a given genomic distance (2 bp to 2001 bp) were calculated by cor.test in R 3.5.0 (https://www.r-project.org/).

### ChIP-seq data processing

Publicly available H3K4me1 ChIP-seq data in mESCs (Feldmann et al., 2020) were downloaded as raw FASTQ files from GEO: GSE141918 with SRA Toolkit v2.11.0 fasterq-dump (http://ncbi.github.io/sra-tools/). All reads were clipped to 35bp in length to excise a region of questionable nucleotide content. Read pairs were then filtered to remove those with mean base quality score less than 20 at either read end. Genomic mapping against the mm10 reference assembly (GRCm38) was performed by Bowtie2 v2.3.0 with parameters “-X 1000 --fr --local -- sensitive-local”. Multiple alignment files per library were combined with Picard tools v1.110 MergeSamFiles.jar, then filtered by samtools v1.3.1 (Li et al., 2009) to retain only proper pair hits with MAPQ score of at least 5.

## Data availability

The datasets generated during this study are available at NCBI Gene Expression Omnibus (GEO) with accession number GSE179266.

## Acknowledgements

We thank Dr. Hua Wang and Dr. Chang Huang in the Helin lab for providing the E14 mESCs and Pa-Tn5 fusion protein, respectively. This work was supported by donations to the Center for Epigenetics Research from The Metropoulos Family Foundation and The Ambrose Monell Foundation and by the Intramural Research Program of the NIH, National Institute of Environmental Health Sciences (ES101965 to P.A.W.).

## Author contributions

R.L. conceived the project and performed the experiments. R.L. and S.A.G. analyzed and interpreted the data. R.L. wrote the manuscript and S.A.G. and P.A.W. reviewed & edited the manuscript.

---

**Figure S1.**
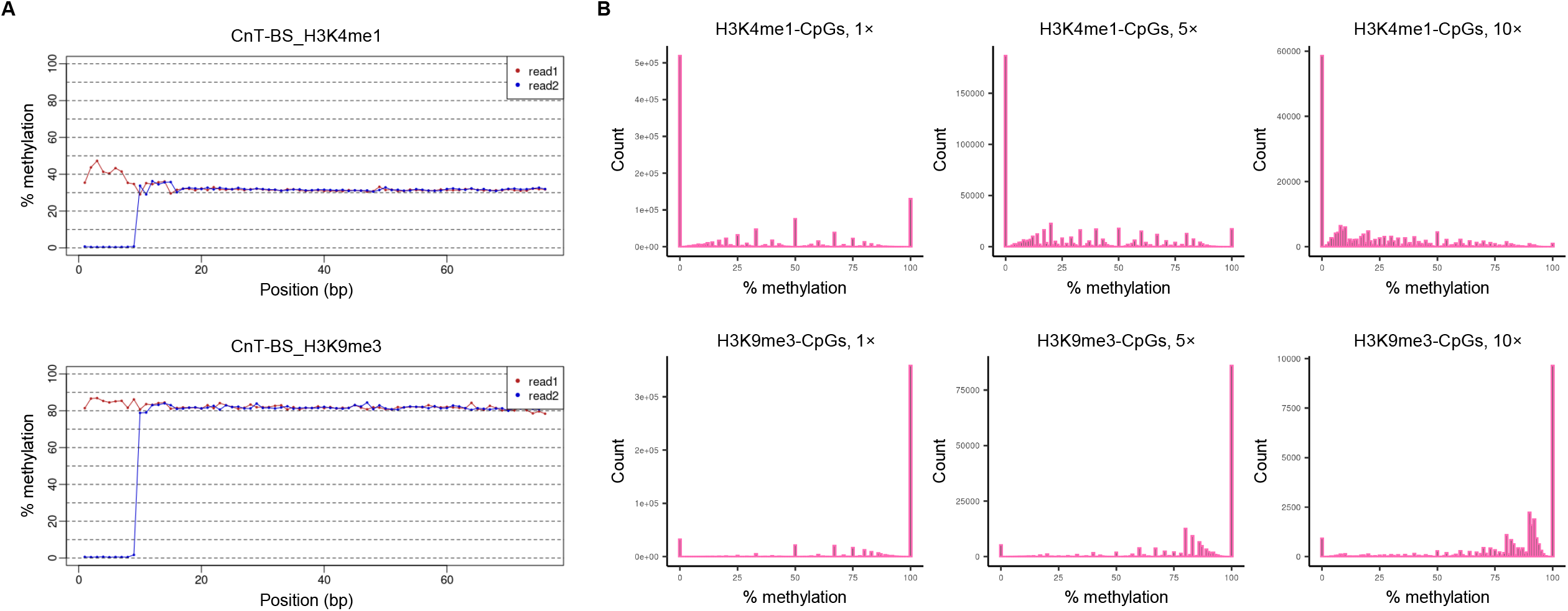
CUT&Tag-BS simultaneously measures DNA methylation, Related to Figure 4. (A) M-bias plots showing averaged CpG methylation rates per position along the reads. (B) Histograms displaying the distribution of methylation levels of individual CpGs at H3K4me1- or H3K9me3-peaks with minimum coverage of 1×, 5×, or 10×.

**Table S1.**
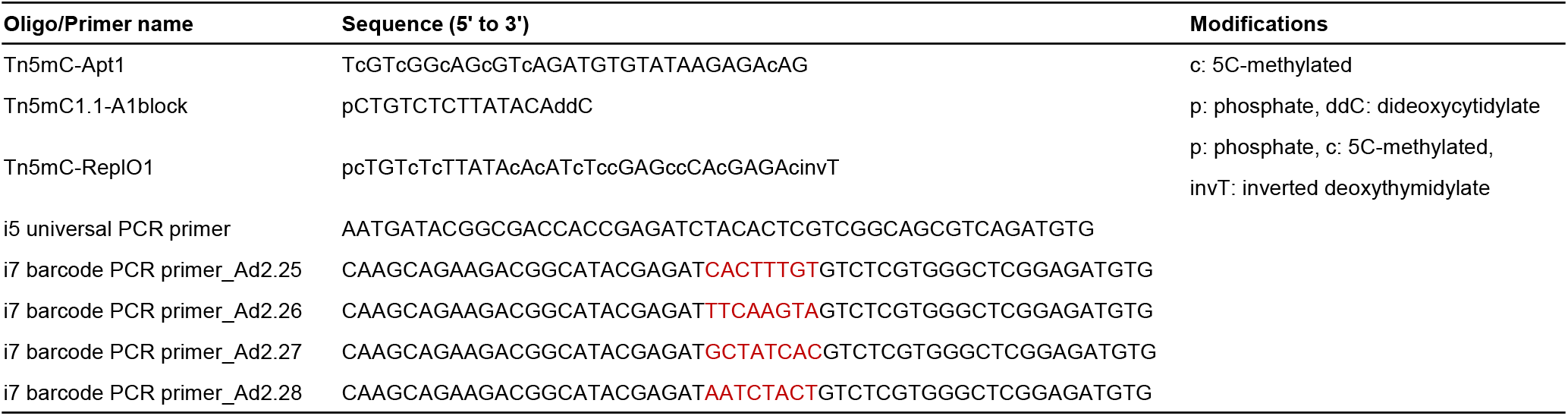
List of oligonucleotides used in this study, related to STAR Methods.

## References

Angermueller, C., Clark, S.J., Lee, H.J., Macaulay, I.C., Teng, M.J., Hu, T.X., Krueger, F., Smallwood, S., Ponting, C.P., Voet, T., et al. (2016). Parallel single-cell sequencing links transcriptional and epigenetic heterogeneity. Nat Methods 13, 229–232.

Bannister, A.J., and Kouzarides, T. (2011). Regulation of chromatin by histone modifications. Cell Res 21, 381–395.

Bell, J.T., Pai, A.A., Pickrell, J.K., Gaffney, D.J., Pique-Regi, R., Degner, J.F., Gilad, Y., and Pritchard, J.K. (2011). DNA methylation patterns associate with genetic and gene expression variation in HapMap cell lines. Genome Biol 12, R10.

Brinkman, A.B., Gu, H., Bartels, S.J., Zhang, Y., Matarese, F., Simmer, F., Marks, H., Bock, C., Gnirke, A., Meissner, A., et al. (2012). Sequential ChIP-bisulfite sequencing enables direct genome-scale investigation of chromatin and DNA methylation cross-talk. Genome Res 22, 1128–1138.

Bulger, M., and Groudine, M. (2011). Functional and mechanistic diversity of distal transcription enhancers. Cell 144, 327–339.

Chen, F., Zhang, Q., Deng, X., Zhang, X., Chen, C., Lv, D., Li, Y., Li, D., Zhang, Y., Li, P., et al. (2018). Conflicts of CpG density and DNA methylation are proximally and distally involved in gene regulation in human and mouse tissues. Epigenetics 13, 721–741.

Clark, S.J., Harrison, J., Paul, C.L., and Frommer, M. (1994). High sensitivity mapping of methylated cytosines. Nucleic Acids Res 22, 2990–2997.

Clark, S.J., Statham, A., Stirzaker, C., Molloy, P.L., and Frommer, M. (2006). DNA methylation: bisulphite modification and analysis. Nat Protoc 1, 2353–2364.

Do, H., and Dobrovic, A. (2015). Sequence artifacts in DNA from formalin-fixed tissues: causes and strategies for minimization. Clin Chem 61, 64–71.

Eckhardt, F., Lewin, J., Cortese, R., Rakyan, V.K., Attwood, J., Burger, M., Burton, J., Cox, T.V., Davies, R., Down, T.A., et al. (2006). DNA methylation profiling of human chromosomes 6, 20 and 22. Nat Genet 38, 1378–1385.

Feldmann, A., Dimitrova, E., Kenney, A., Lastuvkova, A., and Klose, R.J. (2020). CDK-Mediator and FBXL19 prime developmental genes for activation by promoting atypical regulatory interactions. Nucleic Acids Res 48, 2942–2955.

Fu, K., Bonora, G., and Pellegrini, M. (2020). Interactions between core histone marks and DNA methyltransferases predict DNA methylation patterns observed in human cells and tissues. Epigenetics 15, 272–282.

Heinz, S., Benner, C., Spann, N., Bertolino, E., Lin, Y.C., Laslo, P., Cheng, J.X., Murre, C., Singh, H., and Glass, C.K. (2010). Simple combinations of lineage-determining transcription factors prime cis-regulatory elements required for macrophage and B cell identities. Mol Cell 38, 576–589.

Hui, T., Cao, Q., Wegrzyn-Woltosz, J., O’Neill, K., Hammond, C.A., Knapp, D., Laks, E., Moksa, M., Aparicio, S., Eaves, C.J., et al. (2018). High-Resolution Single-Cell DNA Methylation Measurements Reveal Epigenetically Distinct Hematopoietic Stem Cell Subpopulations. Stem Cell Reports 11, 578–592.

Kagey, J.D., Kapoor-Vazirani, P., McCabe, M.T., Powell, D.R., and Vertino, P.M. (2010). Long-term stability of demethylation after transient exposure to 5-aza-2’-deoxycytidine correlates with sustained RNA polymerase II occupancy. Mol Cancer Res 8, 1048–1059.

Kaya-Okur, H.S., Wu, S.J., Codomo, C.A., Pledger, E.S., Bryson, T.D., Henikoff, J.G., Ahmad, K., and Henikoff, S. (2019). CUT&Tag for efficient epigenomic profiling of small samples and single cells. Nat Commun 10, 1930.

Krueger, F., and Andrews, S.R. (2011). Bismark: a flexible aligner and methylation caller for Bisulfite-Seq applications. Bioinformatics 27, 1571–1572.

Langmead, B., and Salzberg, S.L. (2012). Fast gapped-read alignment with Bowtie 2. Nat Methods 9, 357–359.

Lhoumaud, P., Sethia, G., Izzo, F., Sakellaropoulos, T., Snetkova, V., Vidal, S., Badri, S., Cornwell, M., Di Giammartino, D.C., Kim, K.T., et al. (2019). EpiMethylTag: simultaneous detection of ATAC-seq or ChIP-seq signals with DNA methylation. Genome Biol 20, 248.

Li, H., Handsaker, B., Wysoker, A., Fennell, T., Ruan, J., Homer, N., Marth, G., Abecasis, G., Durbin, R., and Genome Project Data Processing, S. (2009). The Sequence Alignment/Map format and SAMtools. Bioinformatics 25, 2078–2079.

Martens, J.H., O’Sullivan, R.J., Braunschweig, U., Opravil, S., Radolf, M., Steinlein, P., and Jenuwein, T. (2005). The profile of repeat-associated histone lysine methylation states in the mouse epigenome. EMBO J 24, 800–812.

Martin, M. (2011). Cutadapt removes adapter sequences from high-throughput sequencing reads. EMBnetjournal 17, 10–12.

Pintacuda, G., Wei, G., Roustan, C., Kirmizitas, B.A., Solcan, N., Cerase, A., Castello, A., Mohammed, S., Moindrot, B., Nesterova, T.B., et al. (2017). hnRNPK Recruits PCGF3/5-PRC1 to the Xist RNA B-Repeat to Establish Polycomb-Mediated Chromosomal Silencing. Mol Cell 68, 955–969 e910.

Quinlan, A.R., and Hall, I.M. (2010). BEDTools: a flexible suite of utilities for comparing genomic features. Bioinformatics 26, 841–842.

Roadmap Epigenomics, C., Kundaje, A., Meuleman, W., Ernst, J., Bilenky, M., Yen, A., Heravi-Moussavi, A., Kheradpour, P., Zhang, Z., Wang, J., et al. (2015). Integrative analysis of 111 reference human epigenomes. Nature 518, 317–330.

Song, Y., van den Berg, P.R., Markoulaki, S., Soldner, F., Dall’Agnese, A., Henninger, J.E., Drotar, J., Rosenau, N., Cohen, M.A., Young, R.A., et al. (2019). Dynamic Enhancer DNA Methylation as Basis for Transcriptional and Cellular Heterogeneity of ESCs. Mol Cell 75, 905–920 e906.

Statham, A.L., Robinson, M.D., Song, J.Z., Coolen, M.W., Stirzaker, C., and Clark, S.J. (2012). Bisulfite sequencing of chromatin immunoprecipitated DNA (BisChIP-seq) directly informs methylation status of histone-modified DNA. Genome Res 22, 1120–1127.

Wang, Q., Gu, L., Adey, A., Radlwimmer, B., Wang, W., Hovestadt, V., Bahr, M., Wolf, S., Shendure, J., Eils, R., et al. (2013). Tagmentation-based whole-genome bisulfite sequencing. Nat Protoc 8, 2022–2032.

Wen, X., Jeong, S., Kim, Y., Bae, J.M., Cho, N.Y., Kim, J.H., and Kang, G.H. (2017). Improved results of LINE-1 methylation analysis in formalin-fixed, paraffin-embedded tissues with the application of a heating step during the DNA extraction process. Clin Epigenetics 9, 1.

Xiao, S., Xie, D., Cao, X., Yu, P., Xing, X., Chen, C.C., Musselman, M., Xie, M., West, F.D., Lewin, H.A., et al. (2012). Comparative epigenomic annotation of regulatory DNA. Cell 149, 1381–1392.

